# Extremely Early Flowering and Large Grain Isogenic Japonica Rice Koshihikari Integrated with Gene *e1* and *GW2*

**DOI:** 10.1101/2025.03.31.646432

**Authors:** Motonori Tomita, Kenta Arai

## Abstract

The extremely early flowering/large-grain isogenic Koshihikari was developed by combining the large-grain allele, *GW2*, derived from Inochinoichi with the year-round flowering allele, *e1*, from Kanto 79. We conducted four back crosses with Koshihikari as a recurrent parent by using an extremely early flowering/large-grain *e1GW2* homozygote as a non-recurrent parent, which was segregated in B_1_F_2_ between Kanto 79 and a large-grain fixed F_3_ plant in Koshihikari × Inochinoichi. In the BC_4_F_2_ population, the *e1GW2* homozygous phenotype was selected and fixed in BC_4_F_3_ as Kshihikari *e1GW2*. plants were segregated according to a ratio of 1 extremely early flowering/large-grain *e1GW2* : 3 extremely early maturing/small-grain *e1gw2* : 3 medium-maturing/large-grain *e1GW2* : 9 medium-flowering/small-grain *E1gw2* and *e1GW2* homozygous isogenic Koshihikari [Koshihiakri *e1GW2*] was fixed in BC_4_F_3_. Whole genome sequencing of Koshihiakri *e1GW2* proved that already known 1 deletion in *GW2* at 8,147,416 bp on chromosome 2, and an *e1* isogenic-specific SNP at 9,090,618 bp, which was 35,213 bp downstream to the 3□ side of *Ghd7* on chromosome 7. RT-qPCR analyses showed that the transcription of *Ghd7* was suppressed in Koshihikari *e1GW2* than in Koshihikari. The Koshihikari *e1GW2* flowered 14 days earlier than Koshihikari and thousand grain weight of Koshihikari *e1GW2* (27.8 g) was 18% larger than Koshihikari (23.6 g). We successfully integrated *GW2* with *e1* for the first time, especially in the genome of a globally produced *Japonica* leading cultivar Koshihikari. The Koshihikari *e1GW2* was registered under plant varietal protection.

## 1. Introduction

Recently, because of the globally aggravated climate crisis, the damage to rice production caused by its lodging is increasing. In Japan, severe and emergency disasters occur frequently; for instance: heavy rains in western Japan in 2018 [1–2], typhoons in eastern Japan in 2019 [3], and heavy rains in July 2020 [4–6]. For example, in 2019, the three large typhoons, Faxai, Tapah, and Hagibis caused paddy rice damage across 45,230 ha, physical damage amounting to 20,320 tons, and monetary damage amounting to approximately 4.9 billion yen [7]. Koshihikari occupies 33.7% of the rice cultivation area and has the highest production volume in Japan [8]. However, Koshishikri has physical defects of long and weak culms and easily falls and is damaged by wind and floods [9]. In addition, the high temperature during the maturation period causes deterioration of rice grain quality, such as taste and appearance [10]. Comparing 50 years before 1970 with 50 years from 1971 to 2021 in Niigata, the highest rice production prefecture in Japan, the average annual temperature increased by approximately 0.8 °C, and the number of times that the annual maximum temperature exceeded 37 °C increased from 5 to 1 5 [11]. Thus, it is no longer feasible to stably produce high-quality rice via open-field cultivation. One of the solutions to avoid unpredictable natural disasters caused by intensifying climate change is to cultivate rice throughout the year regardless of the climate under controlled environmental conditions in a plant factory instead of open-field cultivation. However, the existing rice varieties have a long life cycle, and the cost of plant factory production is 7.1 times that of the paddy field output [12]. Therefore, if the life cycle of rice is shortened to enable year-round production, production costs will decline, and profits will increase.

The authors conducted a genetic analysis using the F_2_, F_3_ of the nonseasonal extremely early flowering mutant Kanto 79, which was induced by γ-ray irradiation of Koshihikari, and the original variety Koshihikari [13]. A single extremely early flowering recessive allele, *e1*,was found to exhibit strong activity resulting in about 14 days earlier flowering in the genetic background of Koshihikari [13]. Moreover, extremely early flowering/semi-dwarf homozygous *e1sd1* and homozygous medium-flowering/semi-dwarf *E1sd1* plants were obtained in Kanto No. 79 (*e1*) × Jukkoku (*sd1*) F_3_; it clarified that *e1* was inherited independently from the semi-dwarf gene *sd1* [13] on chromosome 1 [14–16], which is known to contribute to Green Revolution in rice [17–20]. Then, the medium-flowering *E1sd1* homozygous plant was back-crossed as a non-recurrent plant with Koshihikari eight times to develop Koshihikari *sd1*, known as the variety Hikarishinseiki [13,21–23]. Moreover, back-crossing was conducted five times by using the *e1sd1* homozygous plant fixed in Koshihikari *sd1* × Kanto 79 F_3_ as a non-recurrent plant, with either Koshihikari or Koshihikari *sd1* as the recurrent parent to develop extremely early flowering/semi-dwarf isogenic line Koshihikari *e1sd1*, which was approximately 25 cm semidwarf and flowered 10 days early by Koshihikari[24]. However, extremely early flowering by *e1* reduced yield. Therefore, it is necessary to compensate the yield reduction by combination with any high-yielding genes. In this study, to obtain a short-term growing high-yield rice capable of avoiding severe climatic disasters and high-temperature damage, we developed an extremely early flowering and large-grain isogenic line that combines *e1* and the large-grain gene *GW2* with the genetic background of Koshihikari.

On the other hand, as a result of linkage analysis with DNA markers, the nonseasonalnon-seasonal extremely early flowering gene *e1* was positioned near the flowering suppression gene *Ghd7* (grain number, plant, and heading date 7) localized on the short arm of chromosome 7 [24,25]. Furthermore, whole-genome analysis of the isogenic line Koshihikari *e1sd1*, which was integrated with *e1* in the genetic background of Koshihikari, revealed no mutation in the *Ghd7* gene itself; however, we found a single nucleotide polymorphism (SNP) in the Xa21-like sequence (214 bp) on 35,213 bp downstream to the 3□ side of *Ghd7*that was peculiar to the *e1* line, suggesting its involvement in the suppression of *Ghd7* [24]. *Ghd7* induced by phytochrome receives red light, thereby suppressing the flowering promoting gene [26]. In this study, we conducted a resequencing analysis of Koshihikari *e1GW2* by using the Koshihikari genome as the reference sequence and real-time qPCR analysis on the expression of *Ghd7*, and discussed the relationship between *e1* and *Ghd7*.

## 2. Materials and Methods

### 2.1. Development of extremely early flowering/large-grain Koshihikari *e1gw2*

To combine an extremely early flowering allele, *e1*, from Kanto 79 induced by irradiation to Koshihikari and a large-grain allele, GW,2 from Inochinoichi [27], a large grain plant with *GW2* in F_3_ between Koshihikari and Inochinoichi (Figure 1a) was backcrossed once with Koshihikari, and *GW2* homozygous large grain plant in the BC_1_F_2_ was selected (Figure 1b). Then this plant was crossed with Kanto 79. Since Kanto 79 was regarded to have Koshihikari background, the resulting F_2_ could be considered as BC_2_F_2_. From this BC2F2 population, a homozygous plant showing extremely early flowering and large grain phenotype was selected (Figure 1c) and backcrossed twice with Koshihikari (Figure 1). Finally, in BC_4_F_3_, extremely early flowering and large grain homozygous plants, which were expected to have an *e1GW2* homozygous phenotype, was selected (Figure 4). For *GW2* genotyping, a tightly linked SSR marker, RM3390, was also used. The *e1gw2* homozygous extremely early flowering large-grain plants were selected by measuring the heading date, grain area of all tested plants and genotyping using the DNA marker RM3390 near *GW2* localized at 7.7 Mb from the short arm end of chromosome 2. For the grain area, the grain length and width of each of the three grains were measured for each plant and used as the product of the average values. Genetic analysis was conducted using a trait survey (grain size/ear emergence day) of all plants.

**Figure 1.**
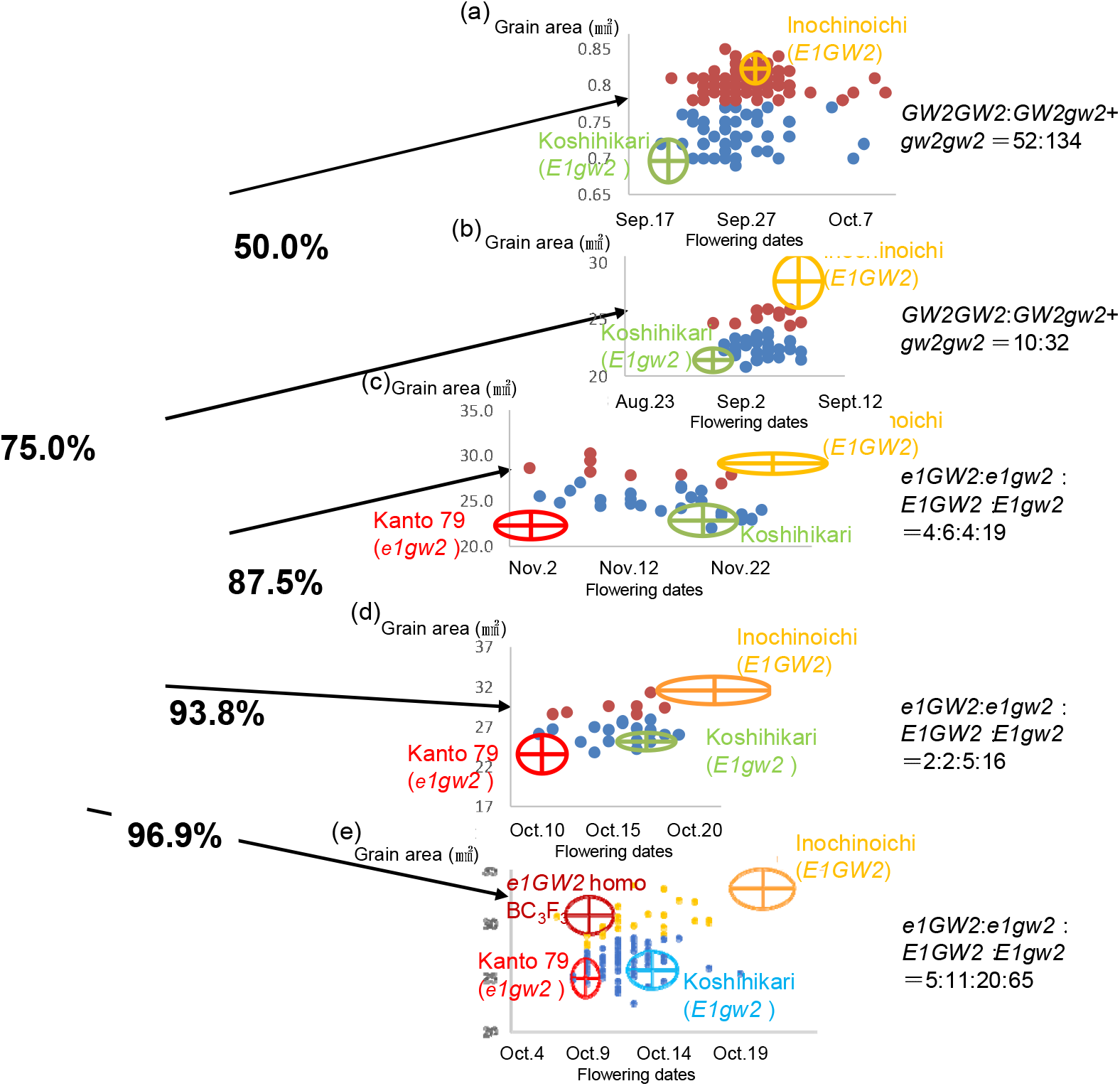
Breeding process of the extremely early flowering large-grain isogenic line Koshihikari e1GW2 To develop an extremely early flowering large-grain line by combining the extremely early flowering gene *e1* of Kanto 79 induced by irradiation to Koshihikari and the large-grain gene *GW2* derived from variety InochinoIchi, we tested B_1_F_2_ to B_4_F_2_ plants according to the cross (Koshihikari*4/[(Koshihikari × InochinoIchi) F_3_ *GW2* type]//Kanto 79), where a *GW2* homozygous large-grain plant fixed in F_3_ between Koshihikari and InochinoIchi was back-crossed four times with Koshihikari.

Cultivation of genetic materials was carried out in a paddy field at Shizuoka University, Shizuoka, Japan, from 2013 to 2021. To accelerate backcross generation, plant materials BCnF_1_ and their next generation BCnF_2_ were grown every year from April to July and July to November, respectively. Namely, we accelerated the generation in a restricted short period. Finally, the acquired genotypes were grown from May to October to test performance. Seedlings were individually transplanted into a paddy field with transplanting density 22.2 seedlings/m^2^ (one seedling per 30 × 15 cm). The paddy field was fertilized by 4.0 kg of basal fertilizer containing nitrogen, phosphorus, and potassium (weight ratio, nitrogen: phosphorus: potassium = 2.6:3.2:2.6) at arate with 4.3 g/m^2^ nitrogen, 5.3 g/m^2^ phosphorus, and 4.3 g/m^2^ potassium across the field. The heading date was recorded as the date first panicle had emerged from the flag leaf sheath for each plant. Culm length was measured as the length between the ground surface and the panicle base. For the yield test, after ripening, ten plants typical of each genotype were sampled twice. The sampled plants were air-dried and were assessed or measured for the following traits, panicle length, number of panicles, number of florets / panicles, proportion of fertile florets, total panicle number, and weight of not milled rice/1,000 grains. Eating tastes were evaluated as seven grades of organoleptic assessment by the panelist and protein contents were determined by Infratec 1241(VOSS Japan Ltd.). The yield of unpolished rice was calculated using the following equation: The yield of not milled rice (g/m^2^) = (number of panicles/m^2^) × (number of florets/panicle) × (proportion of fertile florets) × (weight of not milled rice/grain).

### 2.2. Genetic diagnosis using restriction enzyme digestion with *Hpa I*

PCR was conducted using the dCAPS marker *GW2*-*HpaI* [28], which amplifies 174 bp, including the recognition site for the restriction enzyme *Hpa* I (TaKaRa Bio, Inc., Shiga, Japan) caused by a single nucleotide deletion present in the fourth exon of *GW2*. The composition of the reaction solution with restriction enzyme *Hpa* I was 10 × K buffer (TaKaRa Bio, Inc., Shiga, Japan) 2.5 µL, sterile pure water 12.4 µL, *Hpa* I 0.1 µL, and PCR product 10 µL. The reaction was conducted at 37 °C for 24 h.

### 2.3. Whole genome Sequencing Analysis

We sampled adult leaves of *e1GW2* homozygous plants to extract genomic DNA. Whole genome sequencing was conducted of both Koshishikari *GW2* (BC_6_F_3_) and Koshihkari *e1GW2* (BC_4_F_2_), which was integrated with the extremely early flowering allele, *e1*, and large grain allele *GW2*, by 4 times of backcross into the genetic background of Koshihikari. The leaves were powdered using a mortar and pestle while being frozen in liquid nitrogen. The genomic DNA was then extracted from each cultivar by the CTAB method. Genomic DNA was fragmented and simultaneously tagged so that the peak size of the fragments was approximately 500 bp using the Nextera® transposome (Illumina Inc., San Diego, CA). After purification of the transposome by DNA Clean & ConcentratorTM-5 (Zymo Research, Irvine, CA), adaptor sequences, including the sequencing primers, for fixation on the flow cell were synthesized at both ends of each fragment using polymerase chain reaction, and then the DNA fragments were subjected to size selection using AMPure XP magnetic beads (Beckman Coulter, Brea, CA). Next, the size of the DNA fragment was selected with magnetic beads and qualitatively and quantitatively analyzed to prepare a DNA library. Finally, qualitative checks by using Fragment Analyzer™ (Advanced Analytical Technologies, Heidelberg, Germany) and quantitative measurements by Qubit® 2.0 Fluorometer (Life Technologies; Thermo Fisher Scientific, Inc., Waltham, MA) were performed to prepare a DNA library for NGS. The sequencing was conducted in paired-end 2 × 100 bp on a HiSeq X next-gen sequencer, according to the manufacturer’s protocol (Illumina Inc., San Diego, CA). The gained Illumina reads were firstly trimmed using Trimmomatic (version 0.39) [29] (Figure 2). The sequencing adapters and sequences with low quality scores on 3′ ends (Phred score [Q], <20) were trimmed. The raw Illumina WGS reads were quality checked by performing a quality control with FastQC (version 0.11.9; Babraham Institute). Mapping of reads from Koshihikari *GW2* and Koshishikri *sd1GW2* to the Koshishikri genome as a reference was conducted with Burrows-Wheeler Aligner (BWA) software (version bwa-0.7.17.tar.bz2) [30]. Duplicated reads were removed using Picard (version 2.25.5) (http://broadinstitute.github.io/picard) and secondary aligned reads removed by SAMtools (version 1.10.2) [31]. To identify genetic variations among strains, single nucleotide variant (SNV) detection (variant calling) and SNV matrix generation were performed using GATK < version 4.1.7.0 [32].

**Figure 2.**
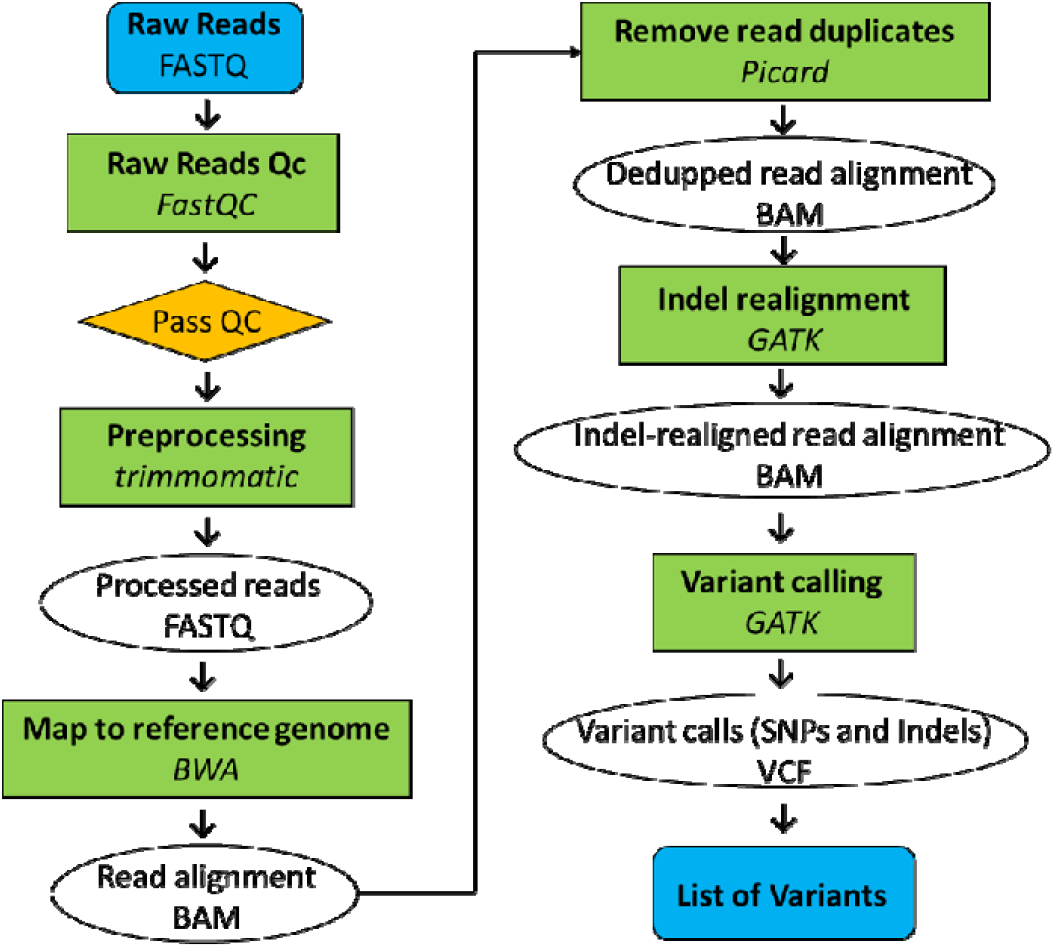
Pipeline of whole genome resequencing analysis.

### 2.4. Genetic diagnosis using restriction enzyme digestion with *Hpa* I

PCR was conducted using the dCAPS marker *GW2*-*HpaI* [28], which amplifies 174 bp, including the recognition site for the restriction enzyme *Hpa* I (TaKaRa Bio, Inc., Shiga, Japan) caused by a single nucleotide deletion present in the fourth exon of *GW2*. The composition of the reaction solution with restriction enzyme *Hpa* I was 10 × K buffer (TaKaRa Bio, Inc., Shiga, Japan) 2.5 µL, sterile pure water 12.4 µL, *Hpa* I 0.1 µL, and PCR product 10 µL. The reaction was conducted at 37 °C for 24 h.

### 2.5. RT-qPCR analysis

Leaves were sampled from Koshishikari and Koshihkari e1GW2 (BC_4_F_3_) before anthesis. Total RNA was extracted from the leaves using the Acid Phenol Guanidium Chloroform method (Chomczynski and Sacchi 2006) and was then treated with 10 U of DNase I (Takara Bio Inc., Kyoto, Japan). poly(A)+ RNA was separated and purified from total RNA using an oligo(dT) cellulose column Poly(A) Quick mRNA Isolation Kit (Stratagene, La Jolla, CA). Single-stranded cDNAs were synthesized by reverse transcription reaction using 0.25 U AMV reverse transcriptase (Life Sciences Advanced Technologies, Saint Petersburg, FL) for 30 min at 50 °C by using 200 ng of poly(A)+RNA as the template of 2.5 *μ*M oligo(dT) primers in 20 *μ*L solution (1 mM dNTP, 5 mM MgCl_2_. 10 mmol/L Tris-HCl (pH 8.3), 50 mmol/L KCl). For RT-qPCR, we designed a FAM-labeled probe from the second exon of *Ghd7*(Os07g0261200), and an 18 mer sense primer (gagcttgaacccaaacacgg) and a 19 mer antisense primer (cataggcttttctggacgcg) that amplified the 118 bp portion of the second exon to sandwich probe. The reaction mixture contained 25 µg of total RNA of Koshishikari or Koshihkari *e1GW2* (BC_4_F_2_), 10 µL of Prime Time Gene Expression Master Mix (2x), and 1 µL of Prime Time qPCR Assay (20x) in a volume of 20 μL. 45 cycles of RT-qPCR comprising denaturation at 95 °C for 15 s and annealing/extension at 60 °C for 45 s were conducted and the fluorescence was detected with the thermal cycler CFX96. Housekeeping gene 18S rRNA [33] was used as a reference for RT-qPCR.

## 3. Results

### 3.1. Inheritance of the extremely early flowering allele, *e1*, and the large-grain allele, *GW2*, in thegenetic background of Koshihikari

In the [Koshihikari/(Koshihikari × InochinoIchi)F_3_ *GW2*]//Kanto No. 79 F_2_, where the large-grain homozygous *GW2* plant that was segregated with the Koshihikari/[Koshihikari × InochinoIchi] F_3_ Gw2 BC_1_F_2_ (grain area 25.8 mm^2^) (Figure 1b) was crossed with Kanto No. 79, a plant group that flowers an average of 13 days earlier and has 28% larger grains than Koshihikari was appeared (Figure 1c). In this study, to accelerate backcross, BC_n_F_1_ and the progeny BC_n_F_2_ were grown every year from April to July and July to November, respectively. Therefore, BC_n_F_2_ grown in a restricted short period tended to distribute continuously in their genetic segregation. Then we estimated genotypes as a reference from the range of parental genotypes and we selected representative phenotypes for the next backcross by the earliest flowering or largest grain in each BC_n_F_2_. The estimated segregation ratios were as follows: extremely early flowering *e1* homozygous :(late-flowering heterozygous *E1e1* + *E1* homozygous) = 10:23, and large-grain homozygous *GW2*:(heterozygous + small-grain *gw2* homozygous) = 8:25; each fitted to a 1:3 ratio. Thus, in a group where the ear emergence day was equivalent to that of Kanto No. 79 (ear emergence day 10/30–11/ 18, 22.1–24.1 mm^2^), the following genotypes were segregated; *e1gw2* homozygous plants (11/10, 30.2 mm^2^) whose the grain area was approximately the same as Inochinoichi (11/19–12/2, 26.7–28.0 mm^2^), and was homozygous for RM3390; *e1e1GW2gw2* plant (11/4–11/11, 24.4–29.4 mm^2^) whose grain area was larger than that of Kanto No. 79 but was heterozygous for RM3390; *e1gw2* type (11/5–11/11, 24.8–27.0 mm^2^), whose grain area was approximately the same as that of Kanto No. 79 and RM3390 was the null type (Figure 1c). On the other hand, in a group whose ear emergence period was approximately the same as that of Koshihikari (11/18–11/24, 21.3–22.7 mm^2^), the following genotypes were segregated; *E1GW2* homozygotes (11/19–11/27, 23.7–27.9 mm^2^), whose grain area was approximately the same as Inochinoichi, and was RM3390 homozygous; *E1GW2gw2* type, whose grain area was between those of Inochinoichi and Koshihikari and was heterozygous for RM3390 (11/14–11/26, 22.9–27.8 mm^2^); and *E1gw2* type (11/14–11/26, 23.0–27.9 mm^2^) with the grain area similar to that of Koshihikari and was the RM3390 null type (Figure 1c). That is, the segregation ratio of BC_2_F_2_ was 4 [*e1GW2* homo]:6 [*e1e1GW2gw2* + *e1gw2* homo]:4 [*E1GW2* homo + *E1e1GW2GW2*]:19 [*E1gw2* homo + *E1E1GW2gw2* + *E1e1gw2gw2* + *E1e1GW2gw2*], which fitted to the theoretical ratio 1:3:3:9 by two genes (χ^2^ = 2.60, 0.45 < P < 0.50) (Figure 1c).

In the Koshihikari///(Koshihikari/[(Koshihikari × Inochinoichi) F_3_ *GW2*]//Kanto No. 79)BC_3_F_2_ (Figure 1d), where an extremely early flowering large-grain homozygous *e1GW2* plant (11/10, grain size 30.6 mm^2^) segregated in B_2_F_2_ (Figure 1c) was underwent a third back-crossing to Koshihikari, an extremely early flowering large-grain *e1GW2* homozygous plant (10/13, 28.6 mm^2^, 10/14, 28.8 mm^2^), early-flowering small-grain *e1gw2* homo, and *e1e1GW2gw2* type plants (10/12, 26.1 mm^2^, 10/13, 26.7 mm^2^) were segregated in the early-flowering group (Figure 1d). Moreover, in the medium-flowering group similar to Koshihikari, large-grain *E1GW2* homo and *E1e1GW2GW2* type plants (10/17–10/20, 28.6–31.3 mm^2^), small-grain *E1gw2* homo, *E1E1GW2gw2, E1e1gw2gw2*, and *E1e1GW2gw2* plants (10/15–10/22, 23.8–27.5 mm^2^) were segregated. That is, the segregation ratio of B_3_F_2_ was 2 [*e1GW2* homo]:2 [*e1gw2* homo + *e1e1GW2gw2*]:5 [*E1GW2* homo + *E1e1GW2GW2*]:16 [*E1gw2* homo + *E1E1GW2gw2* + *E1e1gw2gw2* + *E1e1GW2gw2*], which fitted the theoretical ratio 1:3:3:9 by two-gene segregation (χ^2^ = 3.144, 0.50 < P < 0.55) (Figure 1d). A fourth backcrossing with Koshihikari was conducted on an extremely early flowering large-grain plant (10/14, 28.8 mm^2^) isolated from BC_3_F_2_ (Figure 1c). In the Koshihikari * 2///Koshihikari / [(Koshihikari × InochinoIchi) F_3_ *GW2* type]// Kanto No. 79 BC_4_F_2_, the segregation ratio was 5 extremely early flowering large-grain *e1*Gw2 homozygous:11 extremely early flowering small-grain *e1gw2* type:20 medium-flowering large-grain *E1GW2* type:65 medium-flowering small-grain *E1gw2* type, which fitted to a 1:3:3:9 ratio (χ^2^ = 4.389, 0.15 < P < 0.20) (Figure 1d, 3).

**Figure 3.**
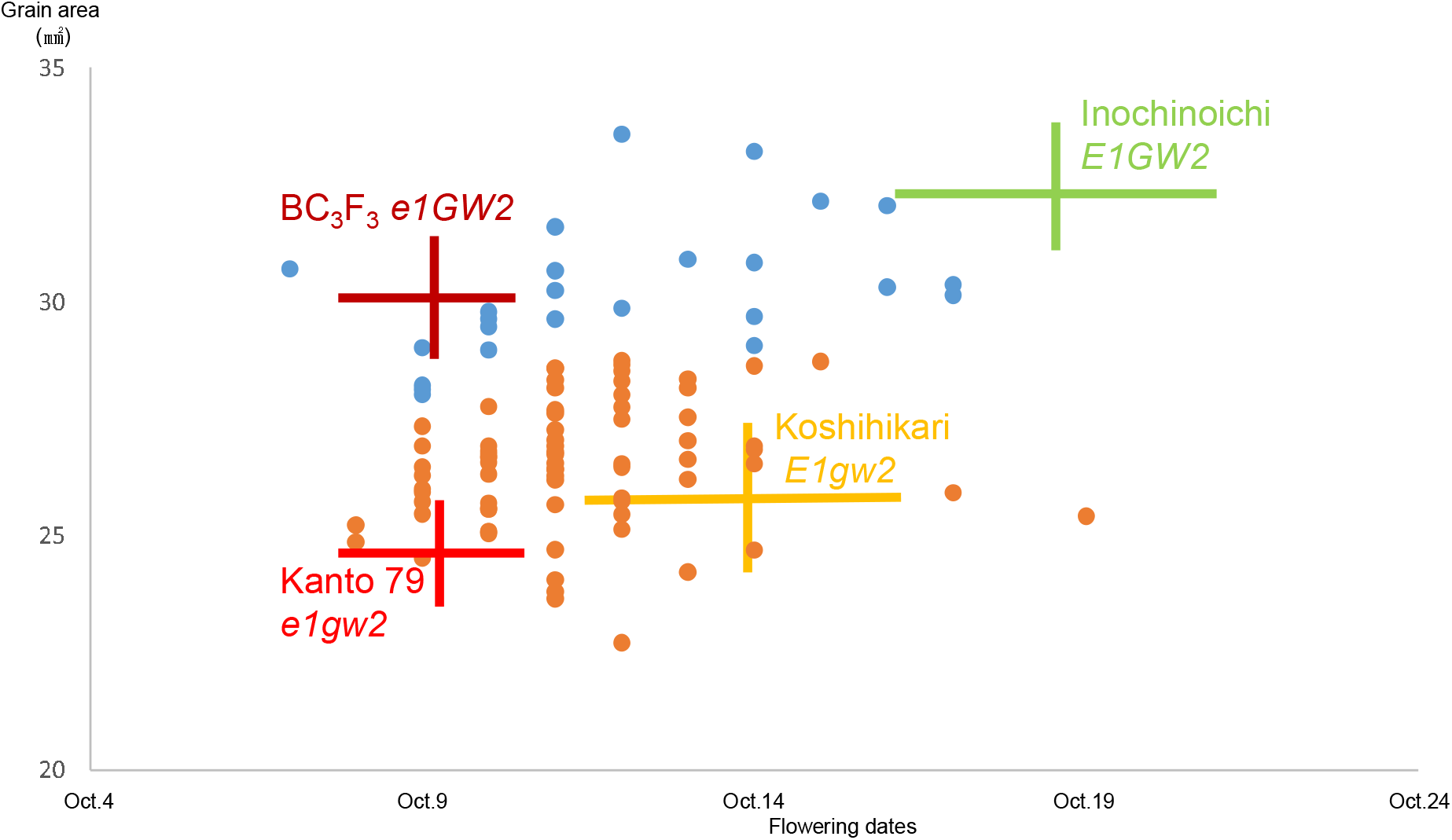
Relationship between ear emergence period and grain area in Koshihikari * 2///Koshihikari / [(Koshihikari × InochinoIchi) F_3_ *GW2*]//Kanto No. 79) BC_4_F_2_.A fourth backcrossing with Koshihikari was conducted on an extremely early flowering large-grain plant (10/14, 28.8 mm^2^) isolated from BC_3_F_2_. In the Koshihikari * 2///Koshihikari / [(Koshihikari × Inochinoichi) F_3_ *GW2* type]// Kanto No. 79 BC_4_F_2_, the segregation ratio was 5 extremely early flowering large-grain *e1*Gw2 homozygous:11 extremely early flowering small-grain *e1gw2* type:20 medium-flowering large-grain *E1*Gw2 type:65 medium-flowering small-grain *E1gw2* type, which fitted to a 1:3:3:9 ratio (χ^2^ = 4.389, 0.15 < P < 0.20).

In the BC_4_F_3_ progenies of of *E1e1GW2gw2* type BC_4_F_2_, regarding the ear emergence day, Kanto No. 79 type extremely early flowering *e1* homozygous and Koshihikari type medium-flowering (*E1* homo + *E1e1* hetero) was segregated into 20:74 ≈ 1:3 (χ^2^ = 0.723) (Figure 4). Regarding grain size, the BC_4_F_3_ progenies were segregated into 27 *GW2* homo:45 *GW2gw2* hetero:22 *gw2* homo being approximately 1:2:1 (χ^2^ = 0.702). Of these, 8 plants were genetically diagnosed by the restriction enzyme *HpaI* digestion. In the plants with *GW2*, the PCR-amplified 174 bp fragment was divided into 151 bp and 23 bp fragments. As a result of genetic diagnosis, 2 of the 8 plants were *GW2* homozygous, 5 were *GW2gw2* heterozygous, and 1 were *gw2* homozygous (Figure 5). The extremely early flowering large-grain homozygous *e1gw2* plant fixed in BC_4_F_3_ had an ear emergence date of September 21, which is approximately the same as that for Kanto 79 and approximately 6 days earlier than that of Koshihikari. Compared to Koshihikari, the panicle length was approximately the same, and the grain area was 29.9 mm^2^, which was 38.6% larger than that of Koshihikari.

**Figure 4.**
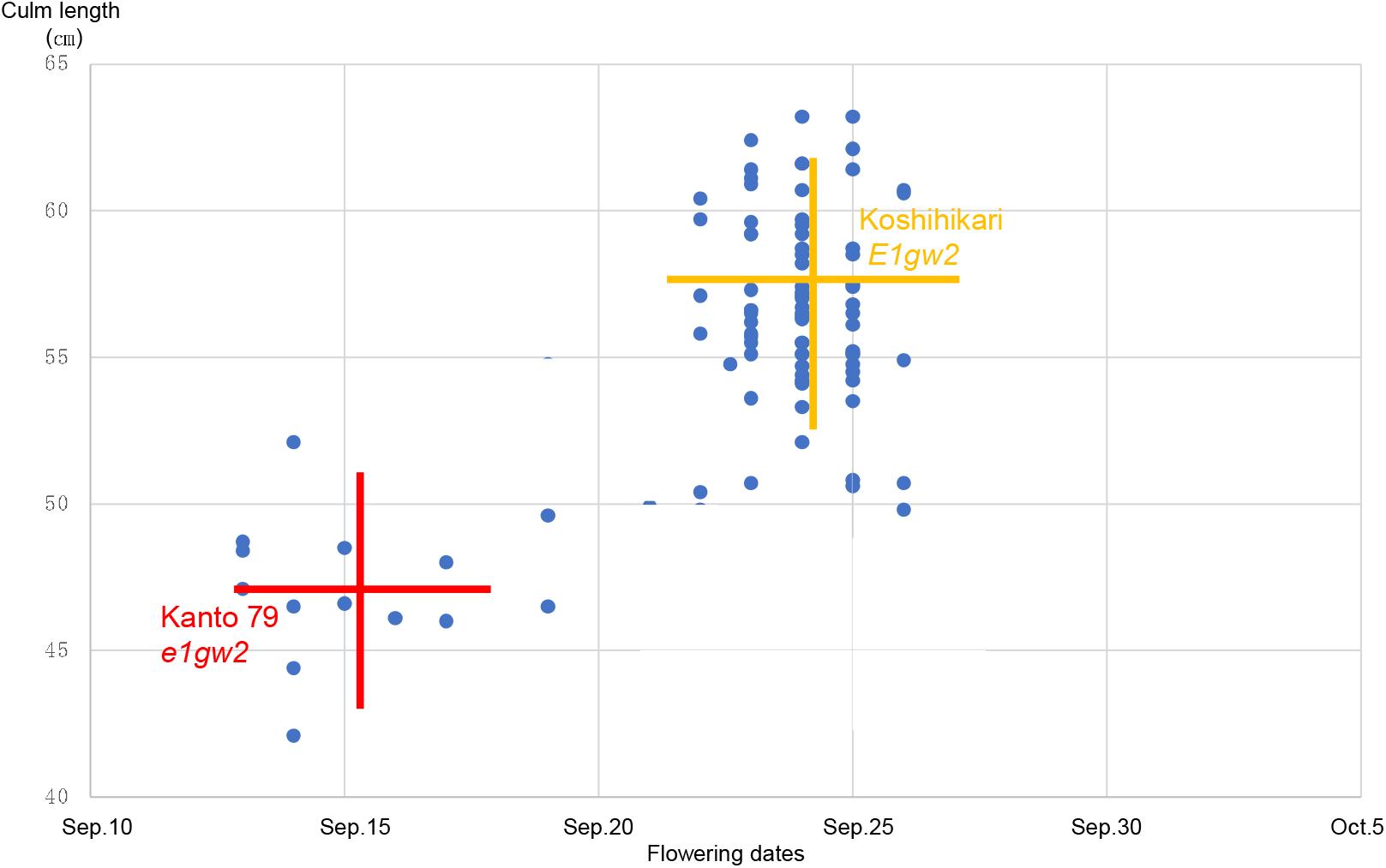
Relationship between ear emergence period and culm length for BC_4_F_3_.In the BC_4_F_3_ progenies of of *E1e1GW2gw2* type BC_4_F_2_, regarding the ear emergence day, Kanto No. 79 type extremely early flowering *e1* homozygous and Koshihikari type medium-flowering (*E1* homo + *E1e1* hetero) was segregated into 20:74 ≈ 1:3 (χ^2^ = 0.723). Regarding grain size, the BC_4_F_3_ progenies were segregated into 27 *GW2* homo:45 Gw2*gw2* hetero:22 *gw2* homo being approximately 1:2:1 (χ^2^ = 0.702).

**Figure 5.**
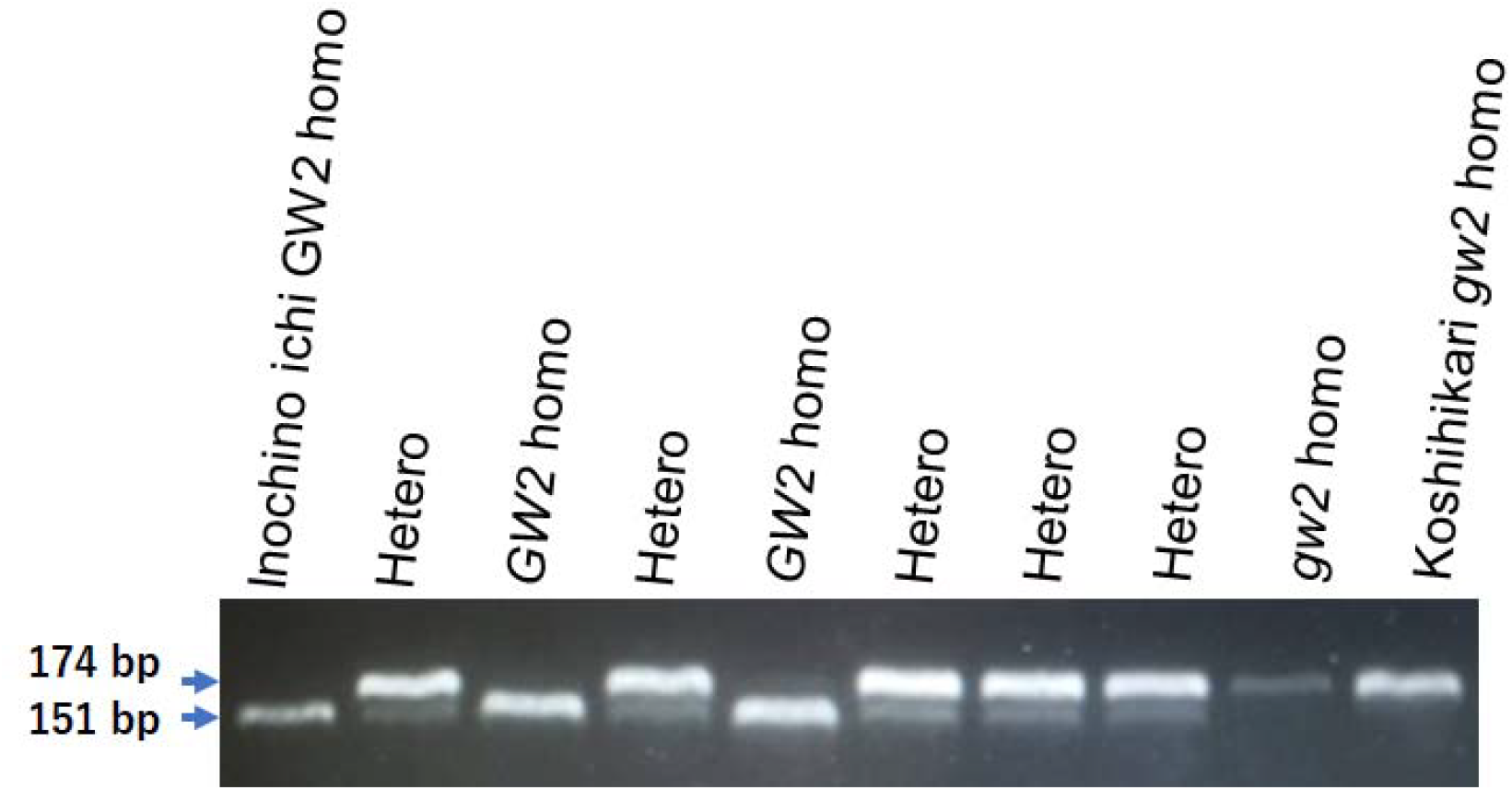
BC_4_F_3_ genetic diagnosis of *GW2* by treatment with restriction enzyme *HpaI*.In BC_4_F_3_, 8 plants were genetically diagnosed by the restriction enzyme *HpaI* digestion. In the plants with *GW2*, the PCR-amplified 174 bp fragment was divided into 151 bp and 23 bp fragments. As a result of genetic diagnosis, 2 of the 8 plants were *GW2* homozygous, 5 were *GW2gw2* heterozygous, and 1 were *gw2* homozygous.

### 3.2. Whole-genome analysis of “Koshihikari *e1GW2*”

The gained reads of Koshishikri *GW2* and Koshishikri *e1GW2* were mapped to the consensus sequence of Koshihikari as the reference, and the mean coverage were 48.41 and 17.55, respectively. As a result of whole-genome analysis of Koshihikari *e1GW2, GW2* was found to be located in the 8,145,094–8,152,058 bp region from the end of the chromosome 2 short arm (single nucleotide deletion at 8,147,417 bp) (Figure 6), and its position was consistent with Koshihikari *GW2*. Except for the region around *GW2*, the number of SNPs were less than 10 per 0.1 Mb. The results indicated that a large portion of the rice 12 chromosomes were substituted with the genome of Koshihikari after continuous backcross. The size of the DNA fragment integrated with *GW2* was determined as the distance between both ends of a SNP cluster (Figure 7). Clusters of SNPs derived from Inochinoichi were distributed over the region of 7,628,443–10,728,113 bp across *GW2*. There existed 166 annotated genes in the integrated DNA fragment. Among them, there were mutations in 2 genes such as hydroquinone glucosyltransferase gene (Table 1). Furthermore, we clarified that Koshihiakri *e1GW2*-specific SNP at 9,090,618 bp, which was 35,213 bp downstream to the 3 □ side of *Ghd7* on chromosome 7 (Figure 8). RT-qPCR results indicated that the rising curve of the fluorescence signal in the *e1GW2* line was slower than that for Koshihikari, and the threshold was reached earlier in the latter (Figure 9). Expression level of 18S rRNA showed the uniformity in relative fluorescence units (RFU) among samples. Therefore, the transcription product *Ghd7* was slightly reduced compared to that of Koshihikari, suggesting that transcription was suppressed in the *e1GW2* line.

**Table 1.** The mutated genes integrated with *GW2* via backcrossing.

| POSITION(bp) | ID | DESCRIPTION |
| --- | --- | --- |
| 7,628,443 | Os02g0234200 | NHL domain-containing protein , Determination of panicle architecture, grain shape and grain weight, FUWA, OsFUWA |
| 8,082,514 | Os02g0242550 | hydroquinone glucosyltransferase |

There existed 166 annotated genes in the integrated DNA fragment. Among them, there were mutations in 2 genes such as hydroquinone glucosyltransferase gene.

**Figure 6.**
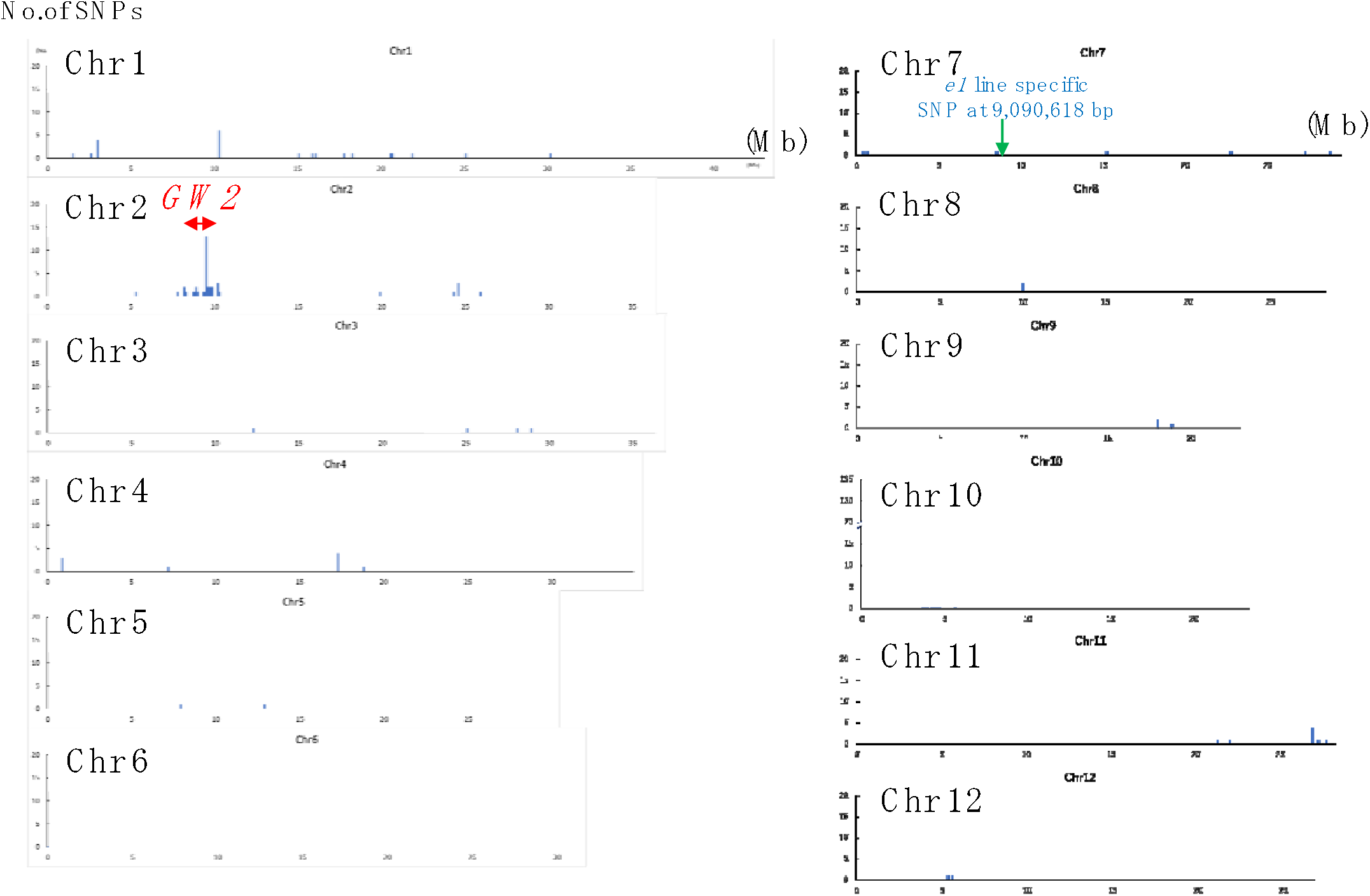
Causative SNP for *GW2* and *e1* in Koshihikari *e1GW2* (BC_4_F_3_). Clusters of SNPs derived from Inochinoichi were distributed over the region of 7,628,443–10,728,113 bp across *GW2*.

**Figure 7.**
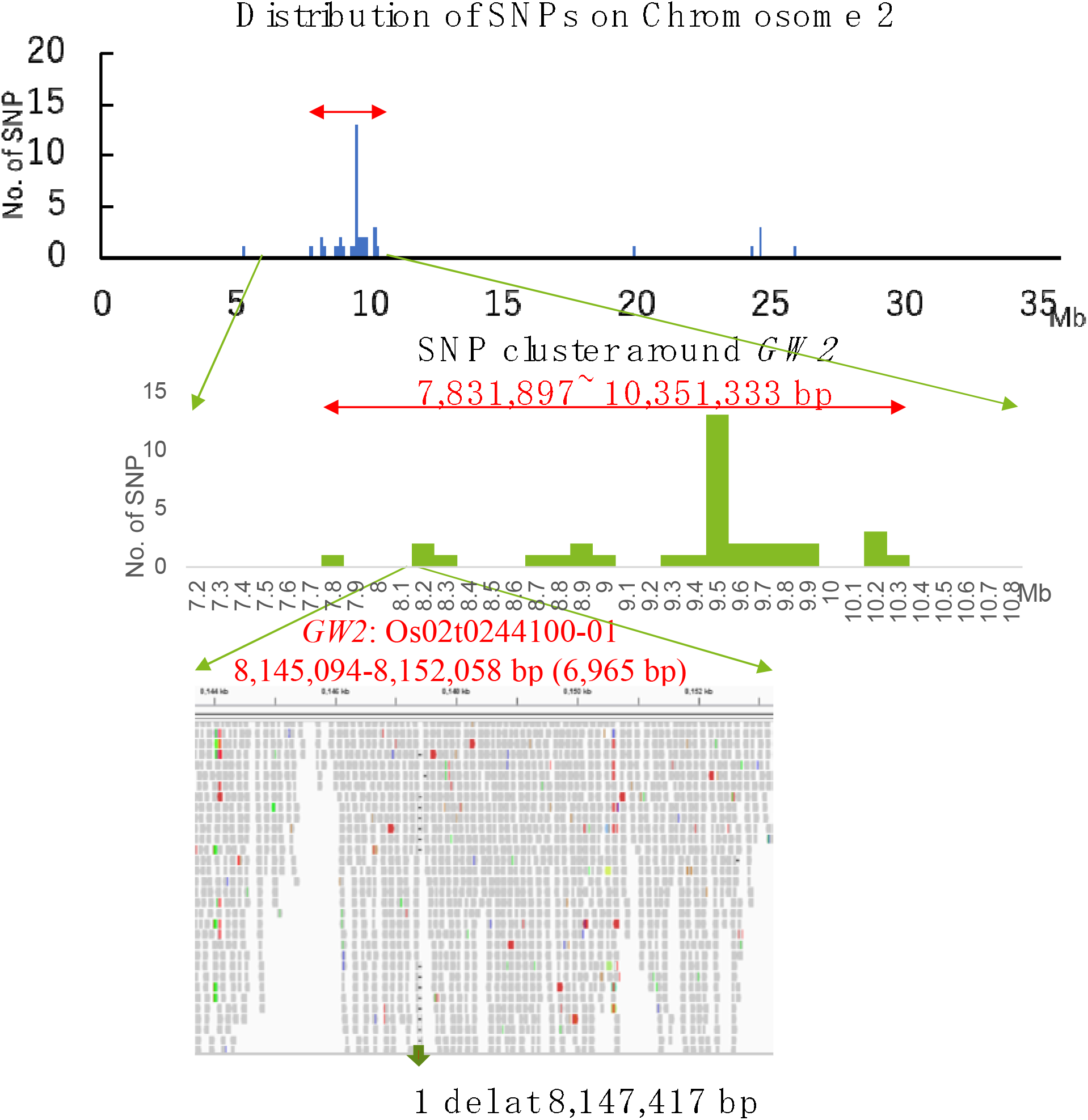
The size of the DNA fragment integrated with GW2. *GW2* was found to be located in the 8,145,094–8,152,058 bp region from the end of the chromosome 2 short arm (single nucleotide deletion at 8,147,417 bp). Except for the region around *GW2*, the number of SNPs were less than 10 per 0.1 Mb. The results indicated that a large portion of the rice 12 chromosomes were substituted with the genome of Koshihikari after continuous backcross.

**Figure 8.**
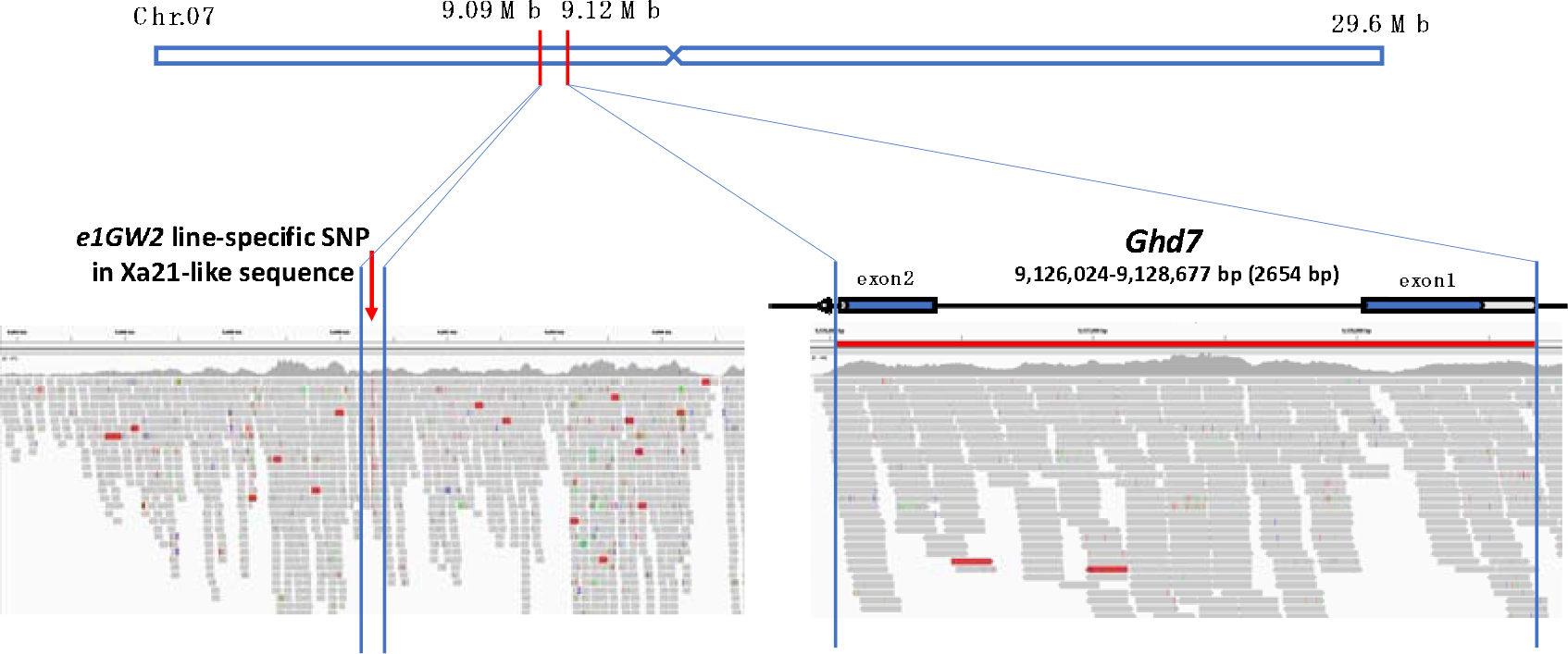
Koshihikari *e1GW2*-specific SNP near *Ghd7*.Whole-genome analysis of the isogenic line Koshihikari *e1GW2*, revealed no mutation in the *Ghd7*and an *e1* line-specific SNP in the Xa21-like sequence (214 bp) on 35,213 bp downstream to the 3’ side of *Ghd7* at 9,090,618 bp on chromosome 7.

**Figure 9.**
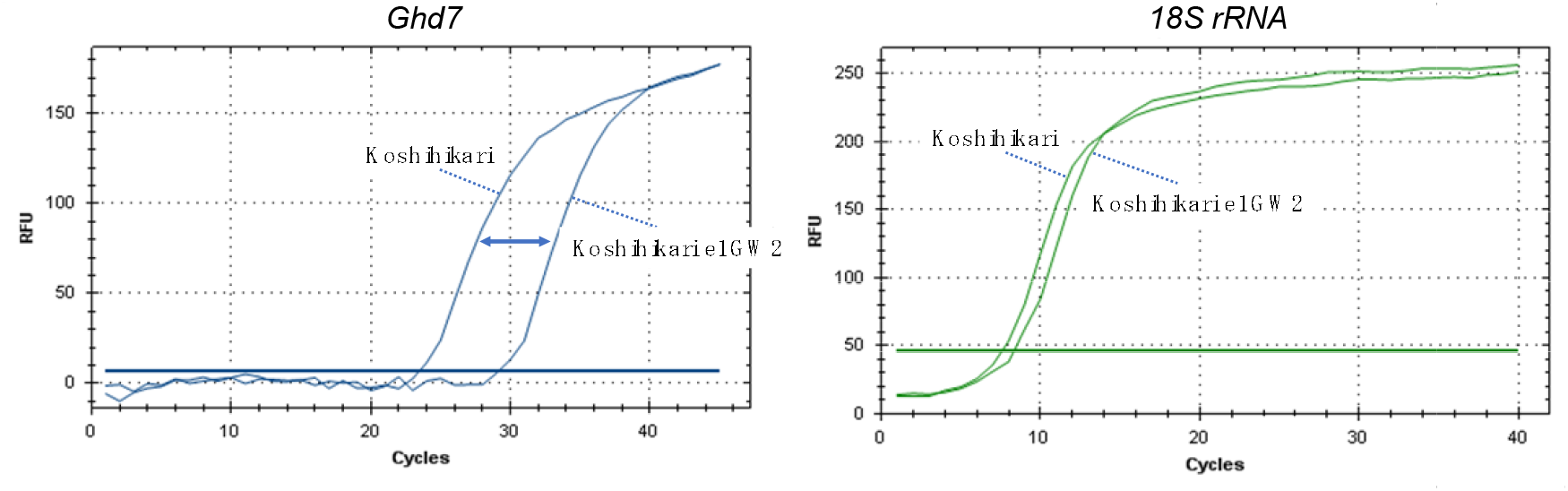
RT-qPCR for *Ghd7*.RT-qPCR results indicated that the rising curve of the fluorescence signal in the *e1GW2* line was slower than that for Koshihikari, and the threshold was reached earlier in the latter. Expression level of 18S rRNA showed the uniformity in relative fluorescence units (RFU) among samples. The result suggested that the transcription of *Ghd7* was slightly suppressed compared with that of Koshihikari.

### 3.3. Trait expression of “Koshihikari *e1GW2*”, an early flowering/large-grain type isogenic line that integrates *e1* and *GW2* in the genetic background of Koshihikari

Compared with Koshihikari, the extremely early flowering/large-grain isogenic line Koshihikari *e1GW2*, was 18.1 cm shorter (Figure 10). We measured the length, width, and surface area of grains on the tip of the main stem and compared these values with those obtained in Koshihikari. We found that grain length and width of Koshihikari *e1GW2* (7.39 mm, 3.84 mm) were approximately 1.08 times and 1.15 times those of Koshihikari (6.85 mm, 3.33 mm), ewspectively. The grain surface area of Koshihikari *e1GW2* (28.4 mm^2^) was 24% larger than that of Koshihikari (22.8 cm^2^) (Figure 9). Thousand grain weight of Koshihikari *e1GW2* (26.0 g) was 10% larger than Koshihikari (23.6 g), and the grain yield of Koshishikari *e1GW2* (42.6 kg/a) was 8% higher than that of Koshihikari *e1* (39.5 kg/a) (Table 2).

**Table 2.** Comparison of agronomic characters of Koshihikari, Koshishikri *GW2*, Koshihikari *e1* and Koshihikari *e1GW2*.

| Genotypes | Days to Heading | Culm length (cm) | Panicle length (cm) | No. of Panicles (No./m<sup>2</sup>) | Grain yield (kg/a) | 1000-grain weight (g) | Protein content | Value of taste |
| --- | --- | --- | --- | --- | --- | --- | --- | --- |
| Koshihikari | 76.8 | 89.6 | 20.0 | 347 | 42.4 | 23.6 | 6.9 | 0.00 |
| Koshihikari GW 2 | 76.3 | 89.2 | 21.8 | 273 | 41.7 | 27.1 | 7.1 | 0.00 |
| Koshihikari e1 | 62.5 | 72.8 | 19.3 | 332 | 39.5 | 22.3 | 6.9 | 0.00 |
| Koshihikari e1GW2 | 62.2 | 71.5 | 18.6 | 386 | 42.6 | 26.0 | 7.0 | 0.01 |

Koshihikari e1GW2
Thousand grain weight of Koshihikari e1GW2 (26.0 g) was 10% larger than Koshihikari (23.6 g), and the grain yield of Koshihikari e1GW2 (42.6 kg/a) was 8% higher than that of Koshihikari e1 (39.5 kg/a).

**Figure 10.**
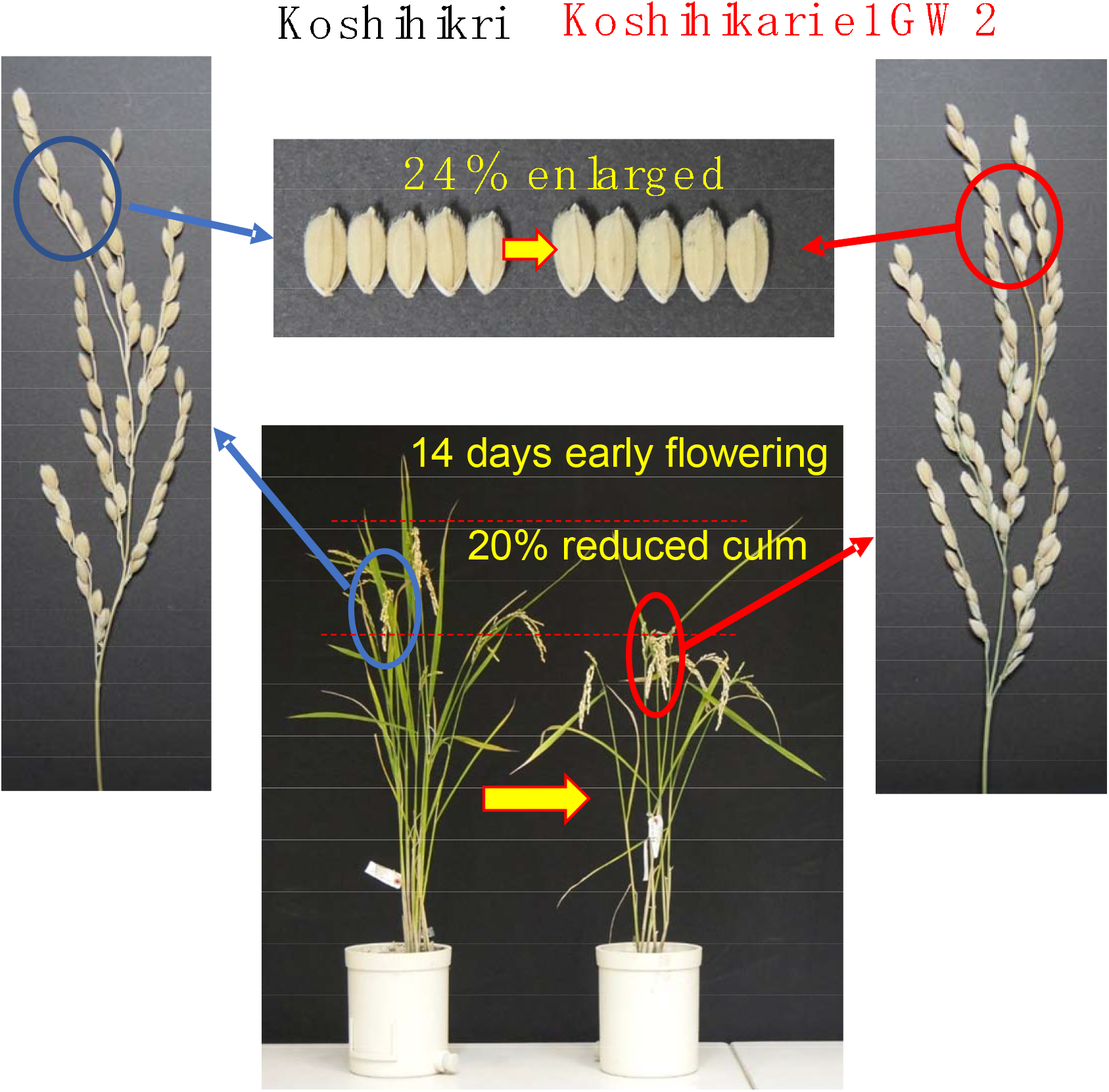
Phenotype of Koshihikari *e1GW2* ([Koshihikari * 2///(Koshihikari / [(Koshihikari × Inochinoichi) F_3_ *GW2* type])//Kanto No. 79] BC_4_F_3_.Compared with Koshihikari, the early flowering/large-grain isogenic line Koshihikari *e1GW2*, which was fixed to the *e1GW2* homozygous genotype in BC_4_F_3,_ was 18.1 cm shorter. The grain surface area of Koshihikari *e1GW2* (28.4 mm^2^) was 24% larger than that of Koshihikari (22.6 cm^2^).

## 4. Discussion

Koshihikari, boasting the largest acreage under rice cultivation in Japan, suffers from and lodging damage as well as high temperatures under climate crisis. In this study, we modified Koshishikari to extremely early flowering by *e1* and further combined the large-grain gene *GW2* to compensate the reduced yield by extremely early flowering, namely. Koshihikari *e1GW2*, that matured 10 days earlier and 10% larger in grain size than Koshihikari. There was no public database on the consensus sequence of Koshihikari, thus, we constructed a consensus sequence of Koshihikari via a high-coveraged whole-genome analysis. We conducted resequencing analyses by using a consensus sequence of Koshihikari as reference. In our study, we clarified that 1 deletion of adenine was detected in *GW2* gene at 8,147,416 bp on chromosome 2, and a SNP in the Xa21-like sequence (214 bp) on 35,213 bp downstream to the 3 □ side of *Ghd7* was detected in at 9,090,618 bp of chromosome 7 in an isogenic large grain Koshihikari integrated with *GW2* and *e1*. The size of the DNA fragment integrated with *GW2* was determined as the distance between both ends of a SNP cluster. From the results of RT-PCR and qRT-PCR, it was inferred that the transcription of *Ghd7* (Koshihikari genome, chr.7: 9,126,024–9,128,677 bp) was suppressed in the *e1* line., and that the G → T SNP commonly seen in Koshihikari *e1sd1* (BC_3_F_2_) and Koshihikari *e1GW2* at 9,090,618 bp of chromosome 7 mighty affect expressing the early flowering trait. This extremely early flowering/large-grain isogenic line was registered as Koshihikari Suruga *e1*Gg in October 2021 (Variety Registration No. 28685) [34]. Koshihikari Suruga *e1*Gg, which compensates the reduced yield caused by extremely early flowering with grain size increase, is considered suitable for gaining high yield in a short period cultivation in a plant factory.

Crops in Japan and the world are damaged by climate change caused by global warming. There has been increased damage from lodging caused by severe weather events like the Western Japan floods and multiple typhoons under the recently intensified climate change. Japanese government put an innovation policy into place to contribute to the world through the development of high-yield crops for a “New Green Revolution.” On the other hand, market liberalization through the Comprehensive and Progressive Agreement for Trans-Pacific Partnership (CPTPP) and negotiations for a Trade Agreement on Goods (TAG), may cause international competition in the rice market, so there is a need for low-cost and high-yielding rice. The rice variety “Inochinoichi” is highly rated at both the consumer and producer levels, but the genes that control its large grain size, approximately 1.5 times that of Koshihikari, have not been elucidated, so the production of “Inochinoichi” is limited to the area around Gifu Prefecture. In the previous study, the authors identified from this unused and buried genetic resource Inochinoichi, the gene responsible for the large grain *GW2* [26], and then by applying the gene to develop a large-grain isogenic of Koshihikari (Koshihikari *GW2*) [26]. Grain size-enlarged Koshihikari has the potential to become advantageously discriminated from US-made Koshihikari. MAFF have registered the large grain isogenic Koshihikari, which was integrated with *GW2*, designated as a new plant variety “Koshihikari Suruga Gg” [35], under Japanese varietal protection. Furthermore, in this study, we developed a large-grain/early flowering isogenic line (Koshihikari *e1GW2*) that incorporates both the *GW2* gene and the extremely early flowering gene *e1*, which we believe an early harvest of enlarged grain in the northern limit or plant factories.

*GW2* was reported as the causative gene involved in grain width in large grain *Japonica* rice WY3. This allele (Os02g024410) encodes a RING protein with E3 ubiquitin ligase activity [36], which acts breakdown in the ubiquitin proteasome pathway [37]. RING-type E3 ubiquitin ligase can control seed development by catalyzing the ubiquitination of expansion-like 1 (EXPLA1), a cell wall-loosening protein that increases cell growth [38]. In contrast, *GW2* derived from WY3 lost function by a frame shift due to a single nucleotide deletion. In a near-isogenic line FAZ1 with *GW2* derived from WY3, the width of the awn was extended by 26.2% because of the increased cell number [36]. In our study, *GW2* from Inochinoichi was also identified as a loss of function by the single nucleotide deletion, increasing grain weight by 34% in the genetic background of Koshihikari. By combining a large-grain gene and an extremely early flowering gene, we have developed a new variety. We believe that this new germplasm, which realizes the early harvest of enlarged grain, will make one of the important contributions to give rise to the “New Green Revolution,” It will be necessary to meet the imminent challenges of population growth, climate change, and increasing globalization.

Including this study, the authors developed a Koshihikari Suruga variety group in which Koshihikari became semidwarf, large-grain, early or late flowering [39]. In each case, promising genes that had not been previously identified were integrated in to Koshishikri genome by continuous five to eight times of backcrossing, and the isogenic line, in which the entire genome was substituted to Koshihikari, was selected and fixed by genomic surveying with next-generation sequencing analysis [24,40,41]. In FY2022, performance or adaptability tests for the recommended variety of Koshihikari Suruga series will be conducted in 10 prefectures. Among them, the extremely early flowering semidwarf variety Koshihikari Suruga *e1sd1* was developed by a combination of *e1* and *sd1* [24], and Koshihikari Suruga *e1*d60 was developed by a combination of *e1* and a novel semidwarf gene d60 differing from *sd1* [42-46]. Both lines were subsequently applied for variety registration and their adaptability and performance will be examined at the Kochi Agricultural Research Center. We think that high-yield breakthrough would be brought by integrating/addition of genes for high-yielding such as large grains and increased biomass on the basis of conventional semidwarfism.

## 5. Conclusions

We successfully integrated *GW2* with *e1* for the first time, especially in the genome of *Japonica* leading cultivar Koshihikari, which is globally produced. Thousand grain weight of Koshihikari *e1GW2* (26.0 g) was 10% larger than Koshihikari (23.6 g), and the grain yield of Koshishikari *e1GW2* (42.6 kg/a) was 8% higher than that of Koshihikari *e1* (39.5 kg/a). It is expected as an early harvest of enlarged grain in the northern limit or plant factories.

## Author Contributions

conceptualization—M.T.; methodology—M.T.; investigation—M.T., K.A.; resources—M.T.; writing—original draft preparation—M.T.; writing—review and editing—M.T.; project administration, M.T.; funding acquisition—M.T. All authors have read and agreed to the published version of the manuscript.

## Data Availability Statement

All the data generated in this study are presented in the main manuscript.

